# Spontaneous peripheral oxygen desaturation and apnea events in mice vary by strain and inspired oxygen level

**DOI:** 10.1101/2025.09.01.673515

**Authors:** Hardik Kalra, Anastasiia Vasileva, Charles R. Jedlicka, Mikhail Vasilyev, Michelle A. Buckman, Zishan Zhang, Brian K. Gehlbach, Junjie Liu, Lara R. DeRuisseau, Mark W. Chapleau, Patrick J. Breheny, Michael H. Tomasson, Melissa L. Bates

**Author notes:** **Corresponding author:** Melissa Bates, PhD, MBA, FAPS. Co-director, Integrative Pathophysiology and Genetics Laboratory, University of Iowa, Departments of Internal Medicine and Pediatrics., **Email:**, Or Michael Tomasson, MD. Co-director, Integrative Pathophysiology and Genetics Laboratory, University of Iowa, Department of Internal Medicine, **Email:**. **Author Contribution:** HK, MHT, MLB, LRD, BKG, AV – Conceptualization and methodology; HK, ZZ, AV, CRJ, PJB, MLB– Data collection and analysis; HK, AV, MV, CRJ, JL, LRD, BKG, PJB, MWC, MHT, MLB – Interpretation of results; HK, CRJ, PJB, MWC, MHT, MLB – Figure preparation; HK, CRJ, MAB, MV, AV, PJB, JL, LRD, BKG, MWC, MHT MLB– Manuscript preparation. Denotes an equal contribution (Michael H. Tomasson, Melissa L. Bates). **Disclosures:** Dr. Bates is the Founder and CEO of LSF Medical Solutions and Dr. Tomasson serves as Chief Medical Officer. Their work at LSF Medical Solutions does not overlap topically with the content of this manuscript. Dr. DeRuisseau is employed by Eli Lilly, a pharmaceutical company; this work was completed prior to her employment at Eli Lilly and the opinions, responsibilities, and obligations related to these endeavors are not in any manner affiliated with Eli Lilly. **New and Noteworthy:** This investigation confirms the finding that mice experience apneas and oxygen desaturations in normoxia and addresses breathing dynamics in the four most commonly studied mouse strains. We observed a reduction of these apneas in hypoxic mice and found that most apneas led to a desaturation. Unexpectedly, most desaturations were not preceded by an apnea. This suggests spontaneous desaturation events are not entirely driven by apnea.

## Abstract

Mouse models of chronic intermittent hypoxia are widely used in research to understand the role of sleep apnea in disease pathogenesis. Mice exposed to periodic reductions in F_I_O_2_ model arterial desaturations observed in humans and recapitulate many comorbidities of sleep apnea. Here, we perform a detailed characterization and confirm reports that mice in room air experience spontaneous, periodic desaturation events. We measured peripheral oxygen saturation in the four mouse strains most commonly used in intermittent hypoxia research (C57BL/6J, CD1, BALB/c, and 129S1) and subjected them to conscious barometric plethysmography to measure oxygen desaturations and apneas simultaneously and took measurements across a range of fractional inspired oxygen (F_I_O_2_). As expected, all strains experienced periodic apneas that were followed by desaturations and decreasing F_I_O_2_ resulted in a reduction of spontaneous apneic events (p = 0.001). Surprisingly, most oxygen desaturations were not preceded by apneas or hypopneas, and mice experienced more desaturations at lower F_I_O_2_ (p < 0.001), despite less frequent apneas. Furthermore, we found strain differences in ventilatory response consistent with prior findings and a novel strain difference in 129S1 mice. These data suggest that spontaneous desaturations are caused not only by apneas and hypopneas but also by other mechanisms, independent of respiration. Our findings provide important context for mouse models of sleep apnea and associated diseases, and future work should explore the extent to which these findings are relevant in humans.

## INTRODUCTION

Chronic intermittent hypoxia (CIH) is widely used in research to model sleep apnea and its contribution to disease, including diabetes, hypertension, cancer, and stroke (1-8). Periodic apneas and arterial oxygen desaturations are typically not observed in conscious, normoxic, healthy humans (9). Mice exposed to CIH model the arterial desaturation effects observed in humans with sleep apnea (10), but some mice experience periodic apneas and arterial oxygen desaturations as part of their native physiological state (11-13). This brings into question the role of these native apneas in normal mouse physiology, whether their frequency is impacted by hypoxia, whether they are seen in the most common strains used in CIH research, and whether they occur in both the light and dark cycles.

For example, Bartolucci, et al. observed a high frequency of central and obstructive apneas in both wild-type and a mouse model of Down syndrome during the light cycle that coincides with rapid eye movement and non-rapid eye movement sleep (14). DeRuisseu et al. observed apneas in conscious mice exposed to normoxia at three ages, and the apneas were elevated in a mouse model of Down syndrome (15). While this and previous studies provide valuable insights, it is crucial to address certain limitations. Periodic arterial desaturations have been detected at 3 months of age, but diminish in number at 6 and 12 months (15). Another study observed numerous arterial desaturations over a 3 hour period in C57BL/6J mice, but these were noted after isoflurane exposure (12). It is possible that these events were an after-effect of isoflurane administration, which suppresses chemoreceptor reflex function (16). Because arterial oxygen saturation and apneas have not been evaluated simultaneously, we cannot conclude that the desaturations and apneas are temporally related. Additionally, the impact of hypoxia, dependency on the light/dark cycle, and whether these observations are strain-specific have not been systematically evaluated.

Differences in the hypoxic ventilatory drive between various strains of mice are well-documented (17-23). Hypoxia stimulates the peripheral chemoreceptors, leading to an increased ventilatory response (24). However, this response can vary between mouse strains. CD1 mice reduce their metabolic demand during hypoxia exposure, whereas C57BL/6J mice switch to ketone metabolism (23). In addition to strain differences, mice exhibit ventilatory differences depending on the light/dark cycle (13, 25). Mice exhibit higher minute ventilation (V_E_) and tidal volume (V_T_) during the dark cycle compared to the light cycle (13, 25). Therefore, it is crucial to investigate if these apneas and desaturations are present in other mouse strains, are influenced by graded inspired oxygen concentration, and occur during both the light and dark cycle.

Using the four most common mouse models in CIH research, we hypothesized that apneas and oxygen desaturations are temporally related and that, as a result of an increased ventilatory drive, hypoxia would reduce the frequency of these events. Further, we hypothesized that apneas and oxygen desaturations would be reduced during the dark cycle.

## METHODS

### Mice

The animal protocol was approved by the University of Iowa Institutional Animal Care and Use Committee (IACUC). Six- to twelve-week-old mice were obtained from The Jackson Laboratory (C57BL/6J #000664, n=14, 5 male; 9 female; BALB/c #000651; 129S1 #002448) and Charles River Laboratories (CD1 Strain #022; all other strains, n=10, 5 male; 5 female). Mice were group-housed in an animal housing facility with access to food (NIH-31 Rodent Diet, Envigo) and water ad libitum, in cages with paper bedding at an ambient temperature of 20-26°C. Mice were subjected to the facility’s standard light (6 AM - 6 PM) and dark (6 PM – 6 AM) cycle.

### Experimental Design

#### Peripheral oxygen saturation

First, to confirm that mice experience periodic desaturations in room air, we performed beat-by-beat pulse oximetry in four isoflurane-naïve female C57BL/6J mice previously retired as breeders. Data were collected in their home cage using a MouseOx collar (Size S-M, MouseOx Plus, Starr Life Sciences Corp) for 20 minutes, as previously described (4). The MouseOx collar sensor was connected to STARR-Link (STARR Life Sciences MouseOx Plus, Oakmont, PA), an analog output module that reports peripheral oxygen saturation at 60 Hz and converts the collar signal to an analog voltage output. The STARR-Link device was interfaced with a PowerLab (ADInstruments, Inc., Colorado, CO, USA) for beat-by-beat oxygen saturation analysis.

Data were continuously recorded using Lab Chart, 8.0 (AD Instruments, Colorado Springs, CO, USA, 1000Hz sampling rate). The day before data collection, mice were anesthetized using 3% isoflurane and maintained at 1.5% to remove neck fur using depilatory cream (Nair™).

#### Identification of relevant mouse strains

We identified mouse strains frequently used in CIH research by conducting a literature review in September 2022. Only primary, English-language, peer-reviewed research articles published between 1991 and 2022 resulting from a Boolean search of “intermittent hypoxia” and “mice” were included. To accommodate the large number of studies returned (n=889), a random number generator was used to select 100 random studies from the results. The top four mouse strains were selected for subsequent experiments.

#### Core Temperature

Prior to microchip implantation for core temperature assessment, mice were anesthetized using 5% isoflurane for 5 minutes, and meloxicam was locally injected in the right lower quadrant of the abdominal area prior to microchip implantation. Temperature microchips (Destron Bio-Thermal LifeChips 985, LifeChip system, Destron Fearing, Airport, TX) were implanted subcutaneously, at the immediate left lower quadrant of the abdominal area to monitor the core temperature during the experiment using the microchip reader (Destron Global PocketReader). Two male CD1 mice were excluded from the following experiments due to post-procedural complications from microchip insertion. The final count for CD1 mice for all the following experiments was 3 male and 5 female mice.

#### Pulse oximetry and plethysmography measurements

Beat-by-beat pulse oximetry data were collected using a recording collar placed on the neck as described earlier. Each mouse was briefly anesthetized using 5% isoflurane and maintained at 1.5 -3% for less than 5 minutes to facilitate placement of the MouseOx collar. Mice were then placed into a whole-body plethysmograph, enabling them full mobility. The measurement order was randomized, and each mouse was tested during both the light and dark cycle. Mice were allowed to rest for at least 24 hours between measurements. On the day of the experiment, mice were acclimated to the lab environment for at least an hour at an ambient temperature of 20-26°C.

A pressure transducer connected to a 0.25 L unrestrained barometric plethysmograph (SCIREQ, model WBP-PC1), gas analyzer (Gemini, CWE Inc.), and spirometer (ADInstruments, Colorado Springs, CO, USA) were calibrated based on the manufacturer’s instructions at the beginning of the experiment. The plethysmograph was connected to paired flow meters (VWR, models 97004-644 and 97004-640), which supplied oxygen and nitrogen (AirGas) to create the experimental F_I_O_2_ gas levels. The flow rate was measured using a spirometer and the gas mixture of oxygen and nitrogen was set to 500 mL/min with an error allowance of 50 mL/min. The gas mixture was allowed to flow freely out of the plethysmograph chamber through an outlet tube. The gas content of the chamber was measured using a fast-responding gas analyzer (Gemini, CWE Inc.) placed separately at both the outlet and inlet ports. Each mouse was allowed to acclimate in the plethysmograph for fifty minutes at 0.21 F_I_O_2_, similar to protocols that use a fixed habituation period (26). After acclimation, 10 minutes of pulse oximetry and breathing measurements were taken at 0.21 F_I_O_2_. The F_I_O_2_ was lowered to 0.15 for 10 minutes and then raised to 0.21 for a 10-minute recovery period. This was repeated for 0.12, 0.09, and 0.30 F_I_O_2_, as shown in Supplemental Figure 1 (26). All F_I_O_2_ levels were accepted ± 0.01. The temperature of the chamber was measured using a glass mercury thermometer, and the core temperature of the mice was monitored throughout the experiment using the implantable microchips. Data from the gas analyzer and plethysmograph pressure transducer were recorded using PowerLab and the LabChart data acquisition software package (ADInstruments, Colorado Springs, CO).

**Figure 1.**
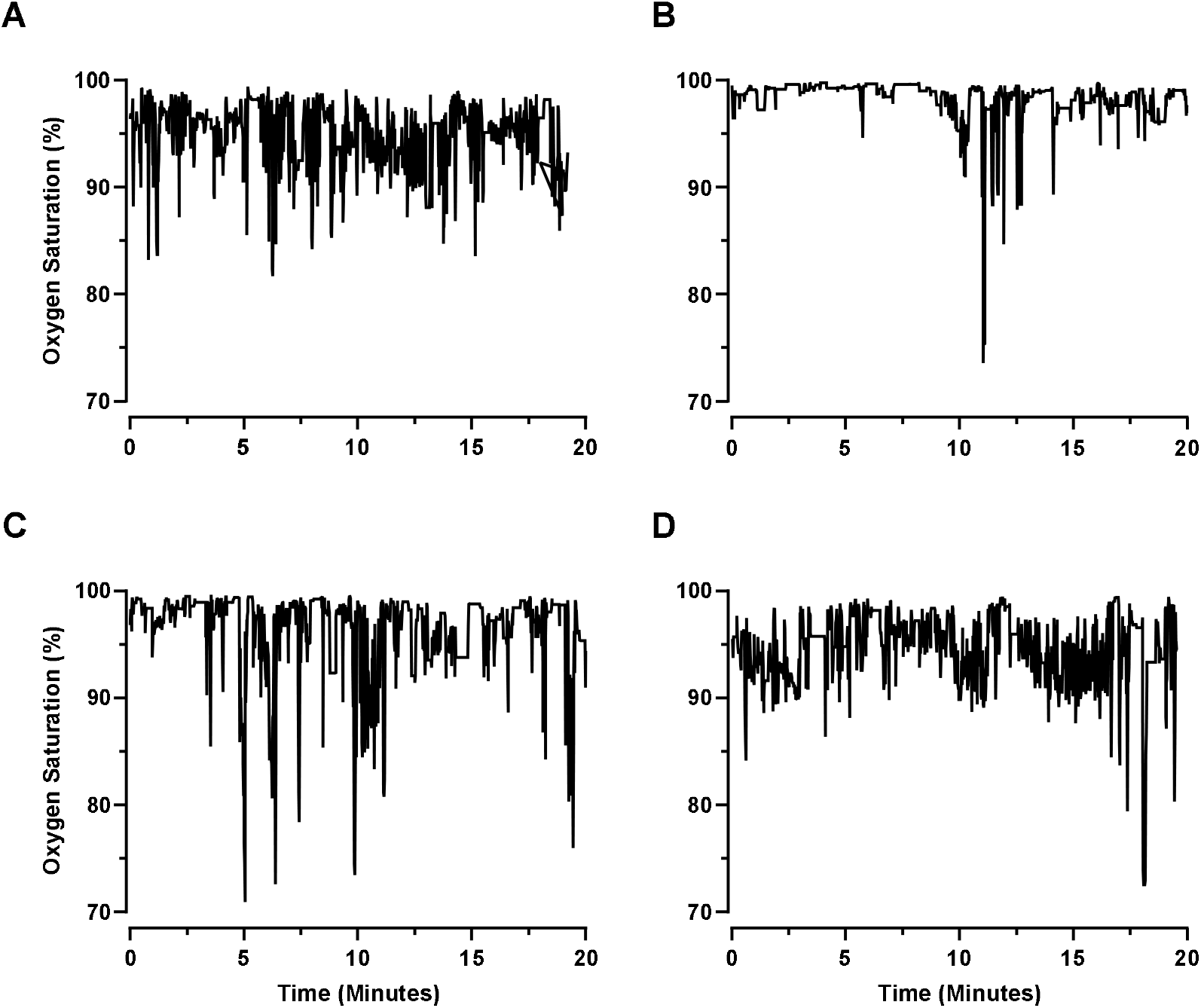
Mean SpO_2_% from measurements of C57BL/6J mice (n=4) in room air for 20 minutes. Spontaneous periodic oxygen desaturations occurred in all animals (7.1 ± 1.59 per minute). Individual mouse data is shown in each panel from A-D.

#### Data analysis

SpO_2_ and breathing waveform data were measured throughout the 10 minutes at each F_I_O_2_ level. Apneas were defined as a cessation of breathing for 0.5 seconds or longer, exceeding the interbreath interval measured during quiet breathing at room air (11). The number and duration of apneas were recorded. Oxygen desaturations were defined based on the American Academy of Sleep Medicine guidelines (AASM) as a decrease in SpO_2_ of 3% or greater from the average during the 10 minutes (27). The number, duration, and severity of desaturations and average SpO_2_ at each F_I_O_2_ level for a 10-minute period were recorded. Desaturation severity was calculated by taking the difference between the average SpO_2_ during the event from the average SpO_2_ over the corresponding 10 minute-period. To determine if apneas and desaturations were associated, we noted each desaturation with an apnea in the preceding 10 seconds. We documented whether there was a desaturation within 10 seconds following an apnea. In rats, the cardiorespiratory response returns to baseline within 10 seconds of an apnea (28), which informed our use of a 10-second window to identify desaturation events. Hypopneas that preceded desaturations were identified and defined as a breathing event with ≥ 30% reduction in tidal volume (29). To associate a hypopnea with a desaturation, we noted each desaturation and searched for hypopneas three seconds prior. The criteria for a hypopnea were informed by the flow tracing from Fleury Curado et al, which shows a flow-limited breath followed by a desaturation within 2-3 seconds (30). Furthermore, we documented whether there was a desaturation within three seconds following a hypopnea. Tidal volume (V_T_), breathing frequency (f_R_), minute ventilation (V_E_), and V_E_/VCO_2_ were expressed relative to average SpO_2_ and calculated using the Drorbaugh and Fenn method, as we have previously (26). V_T_, f_R_, and V_E_ were normalized for body weight to account for size differences between sex and mouse strain. V_E_/VCO_2_ data from the following mice were excluded due to poor data signal quality: two light cycle male BALB/c mice, one dark cycle male BALB/c mouse, and one dark cycle female C57BL/6J mouse.

#### Statistical analysis

Data are represented as mean ± SE, and significance was set *a priori* at p < 0.05. All statistical analyses were conducted using the Minitab software (State College, PA). Two statistical analyses were performed in which the relationship between F_I_O_2_ and other variables was analyzed by repeated-measures analysis of covariance. The first statistical analysis included mouse strain, sex, and light/dark cycle as fixed variables, and F_I_O_2_ as a covariate. Variation in F_I_O_2_ was accepted within ± 0.01. The individual mouse number was included and nested with strain and sex. For the second statistical analysis, F_I_O_2_ × strain, F_I_O_2_ × light/dark, strain × light/dark, sex × strain, and F_I_O_2_ × sex interaction terms were also included. For strain differences relative to average SpO_2_, the strain × SpO_2_ interaction was included. Post hoc analyses were performed using Tukey’s pairwise comparison method, and significance was set *a priori* at p < 0.05.

## RESULTS

### Oxygen desaturations in normoxia

We monitored the peripheral oxygen saturation of four female C57BL/6J mice individually for 20 minutes during the light cycle. We found that all four mice experience periodic oxygen desaturations, as defined by a decrease in SpO_2_ of 3% or greater from the average, while breathing room air (7.1 ± 1.59 per minute, Figure 1).

### Identification of relevant mouse strains

To determine whether the spontaneous hypoxic events we observed were stochastic or influenced by genetic factors, we sought to identify the most common strains used in intermittent hypoxia studies. We found that C57BL/6J mice were used most frequently in intermittent hypoxia studies, followed by BALB/c, 129S1, and CD1 (Figure 2). We utilized these four strains in all subsequent experiments.

**Figure 2.**
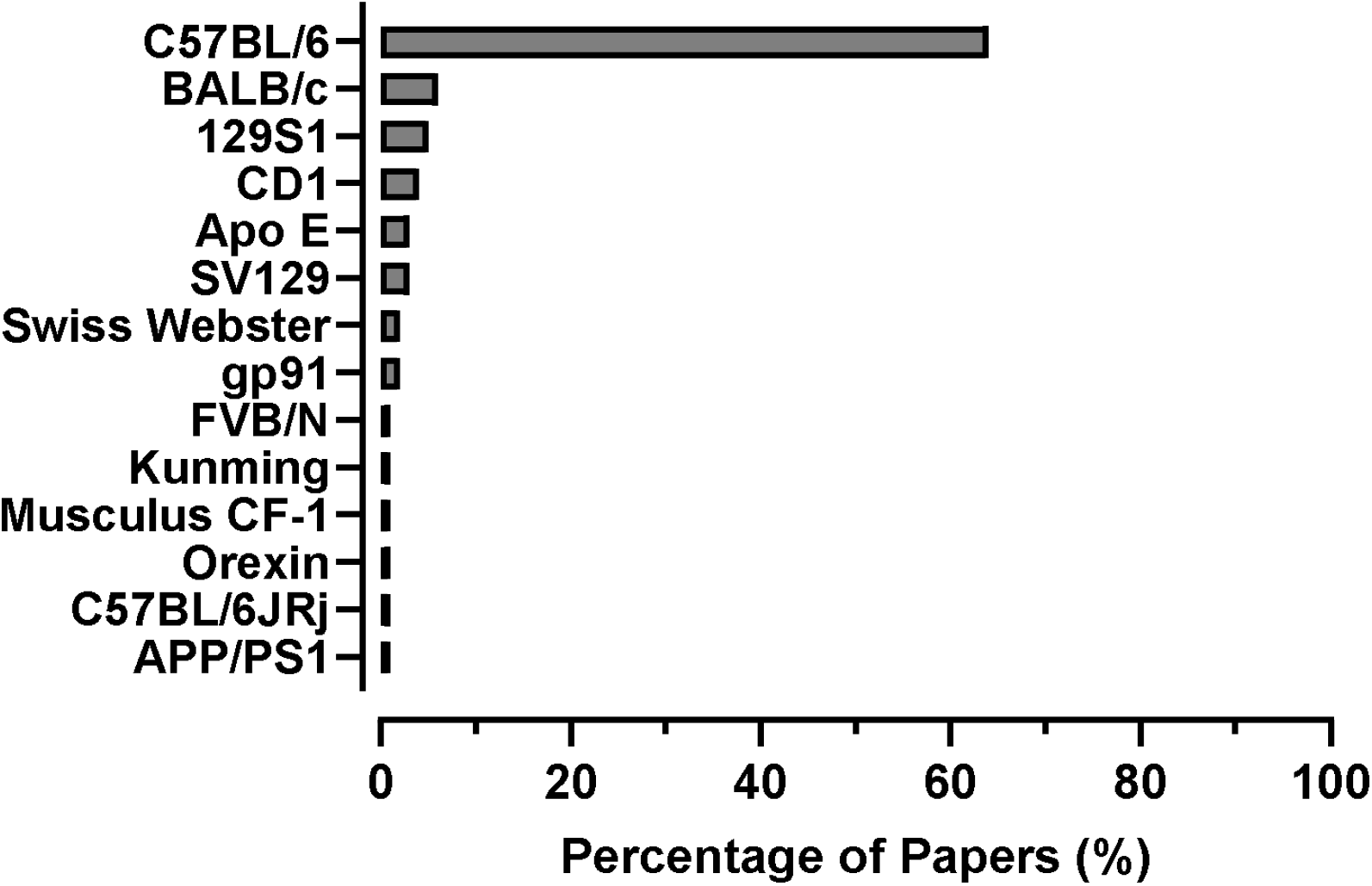
The Web of Science database was used to search for mouse strains used in intermittent hypoxia studies. The relative number of publications (expressed as a percentage) is shown in the graph for 14 strains of mice. The four frequently used mouse strains in intermittent hypoxia research are C57BL/6J, BALB/c, 129S1, and CD1.

### Dose-response relationship between F_I_O_2_ and SpO_2_, apneas, and oxygen desaturations

Control of breathing is a complex physiologic process including a hypoxic respiratory response mediated by the carotid body (31). To assess the degree to which the hypoxic respiratory drive affects SpO_2_, apneas and oxygen desaturations were measured at five levels of F_I_O_2_. As expected, SpO_2_ declined with decreasing F_I_O_2_ (p < 0.001, Figure 3A).

**Figure 3.**
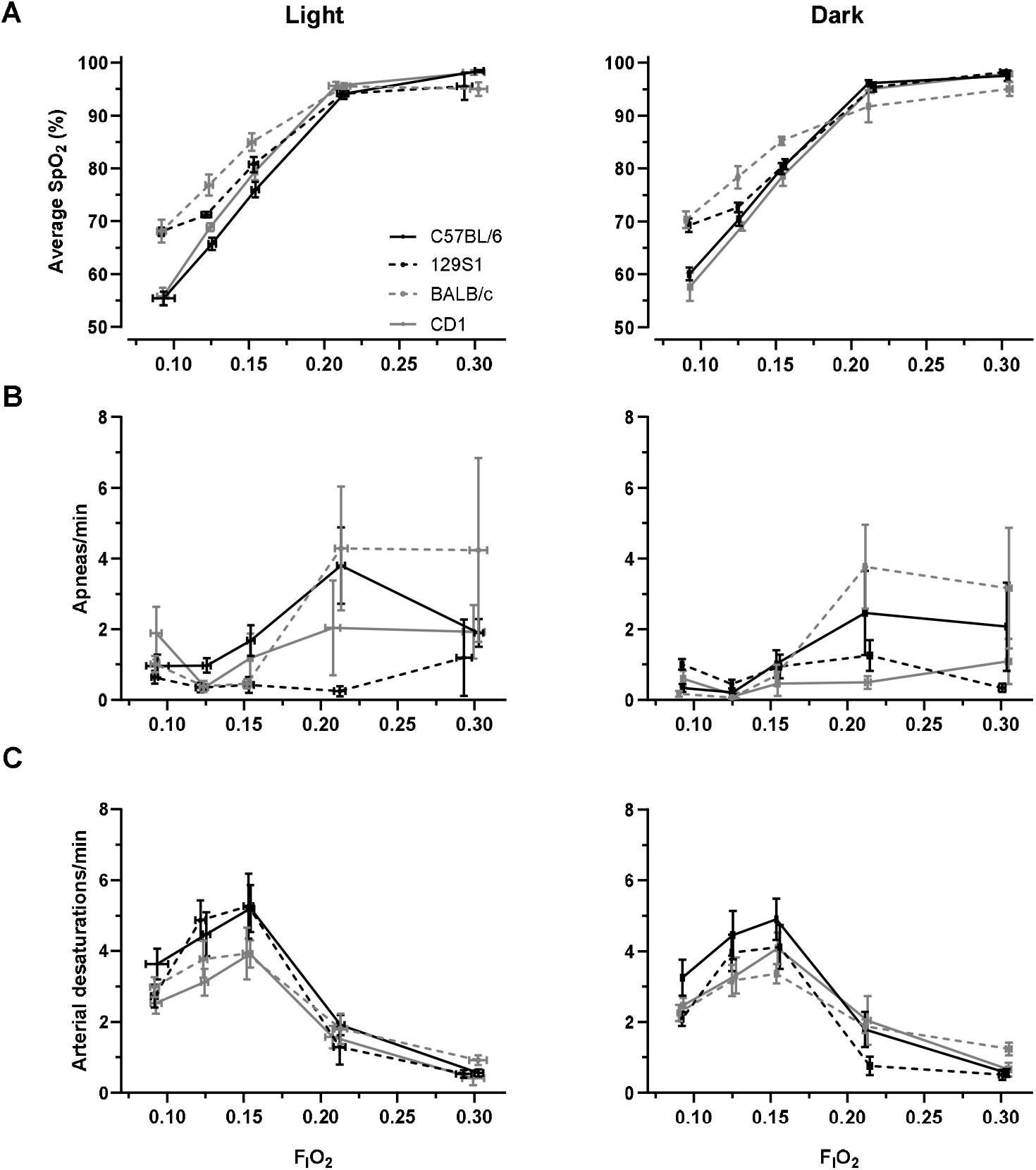
(A) Average SpO_2_ increased with increasing F_I_O_2_ (p < 0.001). F_I_O_2_ × strain differences were observed (p < 0.001). (B) Apneas decreased with decreasing F_I_O_2_ (p < 0.001). F_I_O_2_ × strain was a significant predictor of apneas (p < 0.001). (C) Desaturations increased with decreasing F_I_O_2_ (p < 0.001). F_I_O_2_ × strain was a significant predictor of desaturations (p = 0.040).

The occurrence of apneas declined with decreasing F_I_O_2_ (p = 0.001, Figure 3B). The interaction of F_I_O_2_ and strain was a significant predictor of apneas. Strain differences in the number of apneas were most apparent at higher F_I_O_2_s (p < 0.001, Figure 3B). In a secondary analysis, we found that F_I_O_2_ and sex interaction was a predictor of apneas (p = 0.004). Male mice had fewer apneas than female mice at higher F_I_O_2_.

Contrary to our expectations, mice experienced more desaturations with decreasing F_I_O_2_ (p < 0.001, Figure 3C). Again, F_I_O_2_ and strain interaction was a predictor of the number of oxygen desaturations (p = 0.040). The interaction between F_I_O_2_ and strain was a predictor of SpO_2_ (p < 0.001), demonstrating strain differences in the saturation response to hypoxia. Light/dark cycle did not have an impact on average SpO_2_, number of apneas, and number of desaturations (p = 0.204, 0.067, and 0.128). In a secondary analysis, we found that the interaction of F_I_O_2_ and sex was a predictor of SpO_2_ and desaturation number (p = 0.005). Overall, male mice had a higher SpO_2_ compared to female mice with increasing F_I_O_2_.

### Duration of apnea and oxygen desaturations and severity of desaturations in response to decreasing F_I_O_2_

We found that the duration of apneas increased with decreasing F_I_O_2_ (p = 0.024, Figure 4A) and apneas lasted longer in the light cycle compared to the dark cycle (p = 0.012). The interaction of F_I_O_2_ and sex was a predictor of apnea duration (p = 0.004). The duration of apneas in female mice increased with decreasing F_I_O_2_ compared to male mice.

**Figure 4.**
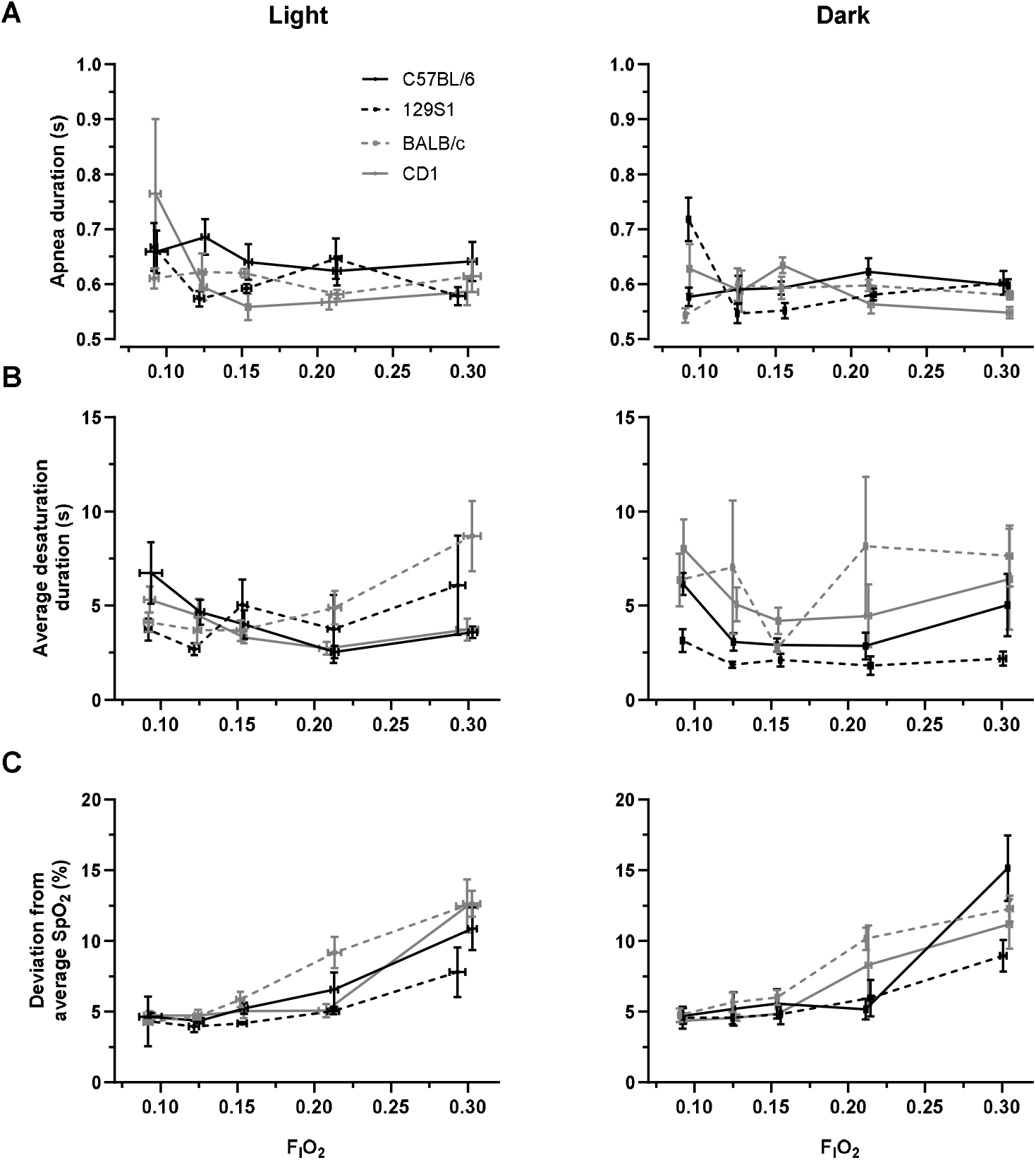
(A) Apnea duration increased with decreasing F_I_O_2_ (p = 0.024). (B) Oxygen desaturation duration did not change with F_I_O_2_ (p = 0.346). (C) Severity of desaturation event increased with increasing F_I_O_2_ (p < 0.001). Strain had no impact on desaturation severity, but F_I_O_2_ and strain interaction was significant (p = 0.214 and 0.003, respectively).

Duration of desaturations did not change with F_I_O_2_ (p = 0.346, Figure 4B). F_I_O_2_ and strain interaction was a predictor of desaturation duration (p = 0.015). Desaturations observed in BALB/c mice lasted longer with increasing F_I_O_2_.

F_I_O_2_ did have a significant impact on the severity of desaturations (p < 0.001, Figure 4C). Desaturations at 0.30 and 0.21 F_I_O_2_ had a 6.62 ± 1.09% and 2.11 ± 0.71% greater drop in SpO_2_ compared to desaturations in hypoxia. The interaction between F_I_O_2_ and strain was a predictor of desaturation severity, with the most severe desaturations observed in BALB/c mice during normoxia (p = 0.003). Light and dark cycles did not impact desaturation severity (p = 0.100). The interaction of F_I_O_2_ and sex impacted desaturation severity (p = 0.032); female mice had an increased desaturation severity compared to male mice at higher F_I_O_2_.

#### Relationship between apnea and desaturation events

To correlate apneas and desaturations, we identified each desaturation and noted any apneas that occurred within 10 seconds prior. The number of desaturations preceded by an apnea was not impacted by decreasing F_I_O_2_ (p = 0.900, Figure 5A). F_I_O_2_ and strain interaction was a significant predictor of desaturations preceded by apneas (p = 0.001), and 129S1 mice and CD1 mice had more desaturations preceded by an apnea with decreasing F_I_O_2_. A strain by light/dark interaction was noted (p < 0.001). Our pairwise comparison showed 129S1 had (16.6 ± 5.97) desaturations preceded by apneas during the dark cycle compared to the light cycle (6.77 ± 2.98, p = 0.046). Whereas, C57BL/6J mice had more desaturations preceded by apneas during the light cycle (21.5 ± 5.67) compared to the dark cycle (8.38 ± 3.00, p = 0.002).

**Figure 5.**
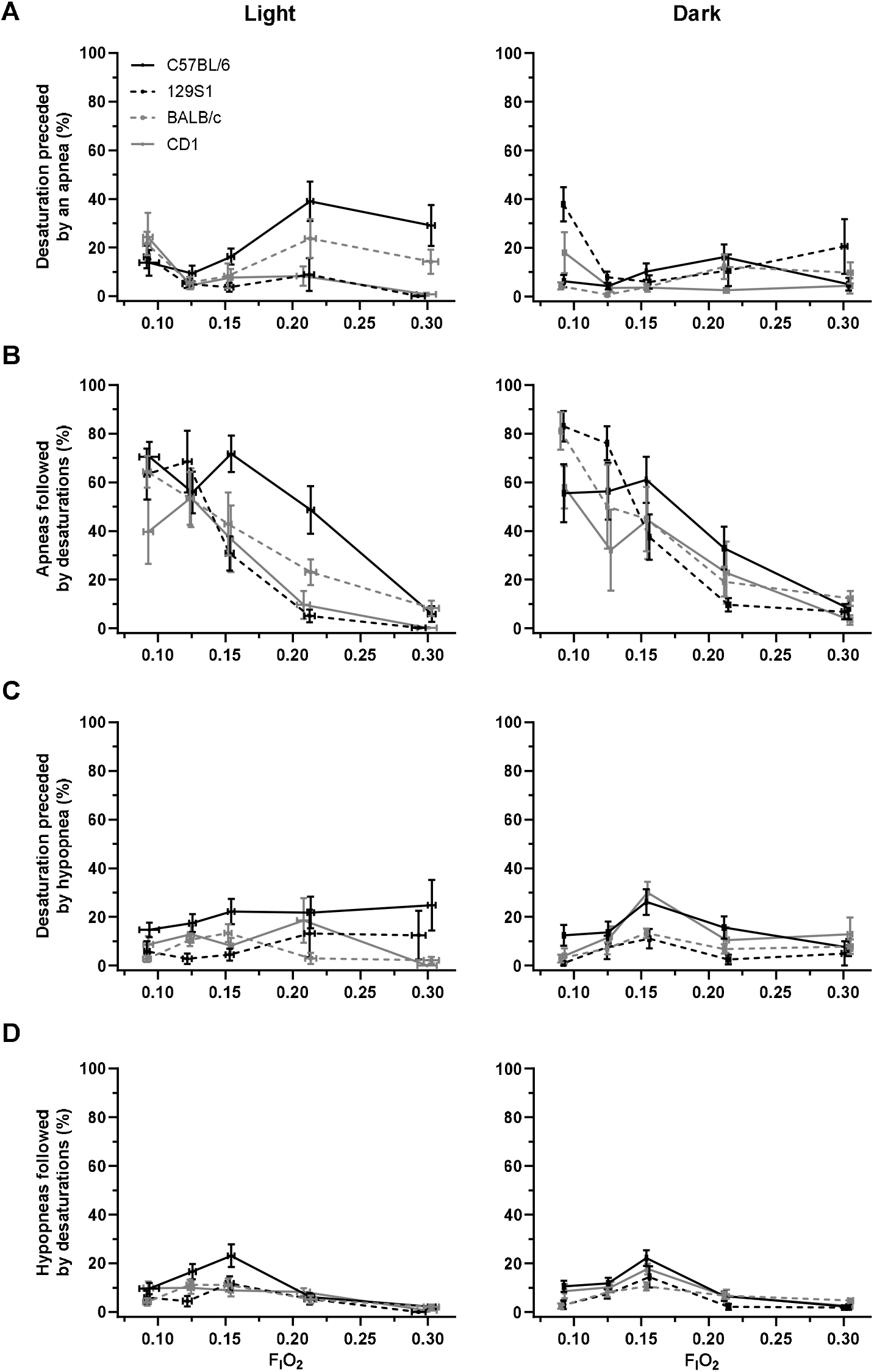
(A) Most desaturations were not preceded by an apnea, and the association decreased further with hypoxia (p = 0.900). Light/dark cycle was a significant predictor of desaturations preceded by apneas (p = 0.026). (B) Apneas were followed by a desaturation with decreasing F_I_O_2_ (p < 0.001). (C) Desaturations were not preceded by a hypopnea (p = 0.739). (D) Hypopneas increased with decreasing F_I_O_2_ (p < 0.001). Furthermore, F_I_O_2_ and strain interaction was a predictor of hypopneas leading to desaturations (p = 0.013). No light/dark or strain differences were observed.

Next, we identified apneas and monitored for a desaturation up to 10 seconds following the apnea to determine whether an apnea preceded a desaturation. Apneas were more often followed by a desaturation with decreasing F_I_O_2_ (p < 0.001, Figure 5B). Strain and light/dark cycle did not have an impact on whether apneas were followed by a desaturation (p = 0.649 and 0.985, respectively).

Notably, desaturations were often not preceded by apneas, with 88% of desaturations occurring independently of an apnea across the strains.

#### Relationship between hypopnea and desaturation events

The number of desaturations preceded by hypopneas was not impacted by decreasing F_I_O_2_ (p = 0.739, Figure 5C). Strain, sex, and light/dark cycle did not impact desaturation and hypopnea association (p = 0.214, 0.860, and 0.604, respectively).

Hypopneas were more often followed by a desaturation with decreasing F_I_O_2_ (p < 0.001, Figure 5D). The interaction between F_I_O_2_ and strain was a significant predictor of hypopneas leading to desaturations (p = 0.013). Sex and light/dark cycle did not impact hypopneas leading to desaturations (p = 0.177 and 0.801, respectively).

#### Ventilatory changes in response to changes in SpO_2_

Overall, a decrease in SpO_2_ did not impact tidal volume (V_T_, p = 0.074, Figure 6A). However, the strain of the mouse was a predictor of V_T_ (p < 0.001). A pairwise comparison revealed that C57BL/6J mice (3.05 ± 0.37 uL/g) had a lower average V_T_ during the stepwise hypoxic challenge compared to 129S1 (4.93 ± 0.70 uL/g, p < 0.001), BALB/c (4.47 ± 0.62 uL/g, p < 0.001), and CD1 mice (4.51 ± 0.41 uL/g, p < 0.001). Furthermore, we observed an increase in strain differences with decreasing SpO_2_ (p = 0.002). 129S1 mice had a higher V_T_ compared to C57BL/6J, BALB/c, and CD1 mice at low SpO_2_. Light/dark cycle was a predictor of V_T_, mice had a higher V_T_ during the dark cycle compared to the light cycle (p < 0.001).

**Figure 6.**
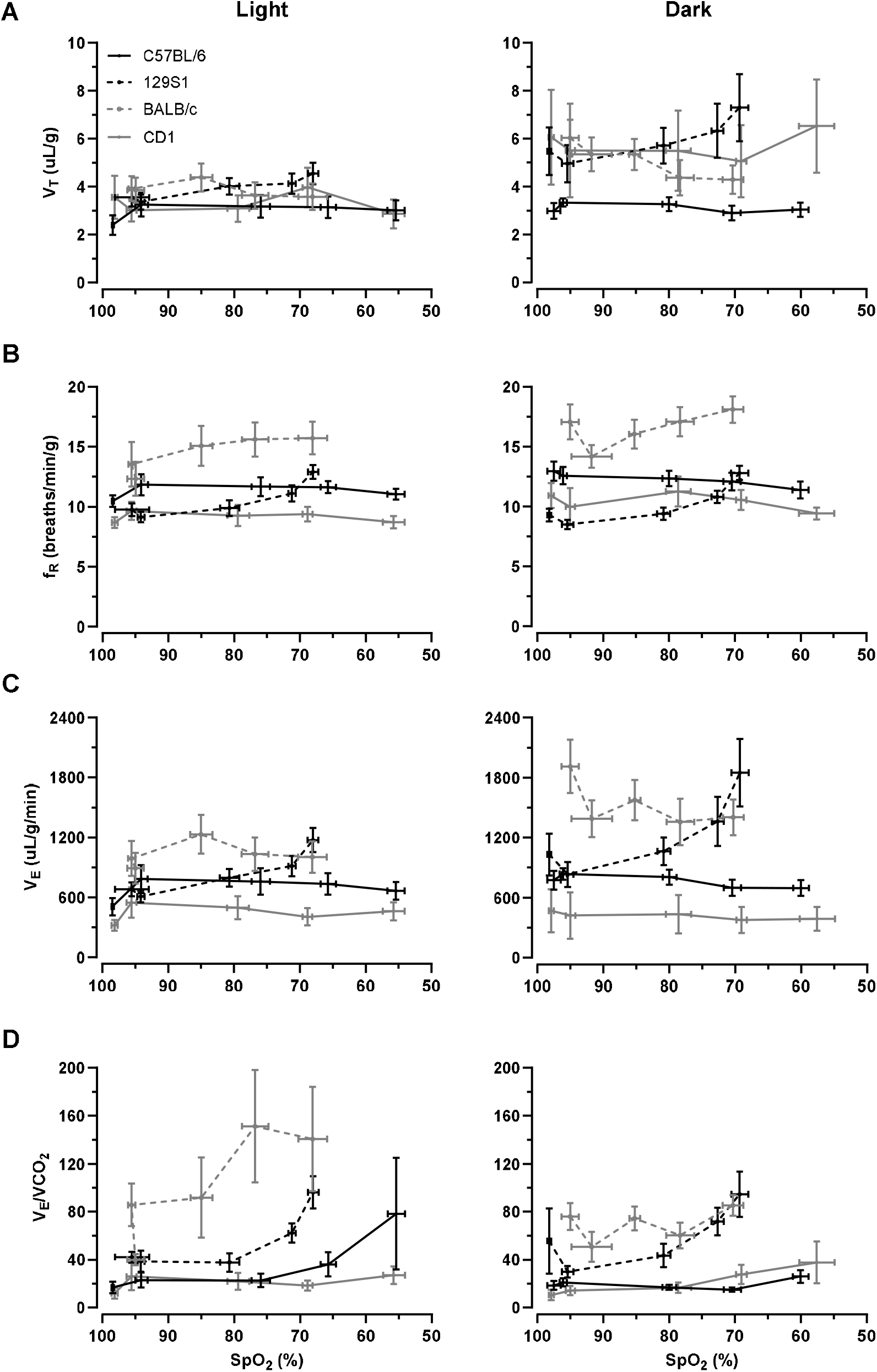
(A) V_T_ (B) f_R_, (C) V_E_, and (D) V_E_/VCO_2_ response to graded F_I_O_2_ plotted vs SpO_2_. Values are shown as means ± SE. (A) V_T_ was not affected by decreasing SpO_2_ (p = 0.074). Strain of the mouse was a significant predictor of V_T_ (p < 0.001). Mice had a higher V_T_ during the dark cycle compared to the light cycle (p < 0.001). (B) f_R_ increased with decreasing SpO_2_. Strain and light/dark were significant predictors of f_R_ (p < 0.001 and 0.001, respectively). (C) V_E_ was not affected by decreasing SpO_2_ (p = 0.156). Strain and light/dark were significant predictors of V_E_ (p < 0.001 and 0.001, respectively). (D) V_E_/VCO_2_ increased with decreasing SpO_2_ (p < 0.001). Strain of the mouse was a significant predictor of V_E_/VCO_2_ (p = 0.001).

Strain and light/dark cycle interaction was observed (p < 0.001). 129S1, BALB/c, and CD1 mice had higher V_T_ during the dark cycle compared to the light cycle (p < 0.001). Although sex did not have an overall impact on the change of V_T_ during a stepwise hypoxic challenge (p = 0.803), sex and strain interaction was a predictor of V_T_ (p < 0.001). V_T_ of 129S1 female mice was (2.59 ± 0.32 uL/g) higher than 129S1 male mice, and (2.35 ± 0.32 uL/g) higher in female BALB/c mice than in male BALB/c mice. Contrastingly, V_T_ of CD1 male mice was (5.04 ± 0.37 uL/g) higher than CD1 female mice.

Reduction in SpO_2_ increased breathing frequency (f_R_, p < 0.001, Figure 6B). Strain of the mouse was an overall predictor of f_R_ (p < 0.001). BALB/c mice had an average of 15.49 ± 1.36 breaths/min/g, C57BL/6J had 11.79 ± 0.67 breaths/min/g, 129S1 mice had 10.35 ± 0.54 breaths/min/g, and CD1 mice had 9.78 ± 0.83 breaths/min/g. There was a significant interaction between SpO_2_ and f_R_ (p < 0.001). 129S1 and BALB/c mice had a greater increase in f_R_ with decreasing SpO_2_ compared to C57BL/6J and CD1 mice. Light/dark cycle impacted f_R_ (p < 0.001), f_R_ was 0.99 ± 0.20 breaths/min/g greater during the dark cycle compared to the light cycle. Sex of the mouse had an impact on breathing frequency (p < 0.001), with 4.07 ± 0.90 more breaths/min/g in female mice compared to male mice.

Overall, the change in SpO_2_ did not impact minute ventilation (V_E_, p = 0.156, Figure 6C), but strain was a significant predictor of V_E_. V_E_ was 1030.47 ± 150.0 uL/g/min in 129S1, 725.29 ± 94.63 uL/g/min in C57BL/6J, 1277.69 ± 190.82 in BALB/c, and 1149.82 ± 419.34 uL/g/min in CD1 mice. A SpO_2_ and strain interaction was observed (p = 0.001). 129S1 mice had a greater increase in V_E_ in response to decreases in SpO_2_ compared to C57BL/6J, CD1, and BALB/c mice. These strain differences were further impacted by light/dark cycle (p < 0.001), 129S1, BALB/c, and CD1 mice had a higher V_E_ during the dark cycle compared to the light. 129S1 and BALB/c female mice had a higher V_E_ compared to their male counterparts, while CD1 male mice had a higher V_E_ compared to CD1 female mice.

Unlike humans, mice vary their metabolic rate in different conditions, e.g., hibernation. We monitored for metabolic rate with the volume of CO_2_ produced per minute (VCO_2_) and found that V_E_/VCO_2_ increased with decreasing SpO_2_ (p < 0.001, Figure 6D). Strain of the mouse impacted V_E_/VCO_2_ (p = 0.001). We found that BALB/c mice had a higher V_E_/VCO_2_ compared to 129S1, CD1, and C57BL/6J mice. Although light/dark cycle did not impact V_E_/VCO_2_ (p = 0.664), strain and light/dark interaction were observed (p < 0.001). CD1 mice had a higher V_E_/VCO_2_ during the dark cycle compared to the light cycle. No sex differences were observed in response to V_E_/VCO_2_ (p = 0.647).

## DISCUSSION

With increasing use of mouse CIH models to study the physiological impacts of sleep apnea, we sought to better understand the relevance and underlying physiology of spontaneous desaturations in mice. We demonstrate spontaneous oxygen desaturations in mice in room air. With the increased respiratory drive upon hypoxia exposure, we found decreased apneas as expected.

However, exposure to hypoxia led to more oxygen desaturations, not fewer. Furthermore, most desaturations could not be explained by apneas, suggesting that mice have an alternative contributor to periodic hypoxemia (32).

### Physiological response to hypoxia

Consistent with previous literature, we observed a strain-dependent response to hypoxia (21-23). BALB/c and 129S1 mice had a higher SpO_2_ during hypoxia compared to C57BL/6J and CD1 mice. A similar difference in SpO_2_ between BALB/c and C57BL/6J mice exposed to hypobaric hypoxia has been shown (21). This difference is especially profound at lower F_I_O_2,_ as demonstrated by the interaction between F_I_O_2_ and strain (p < 0.001). A plausible reason for the higher SpO_2_ observed during hypoxia in BALB/c mice may be due to enhanced hypoxic pulmonary vasoconstriction compared to C57BL/6J mice that more efficiently maintain ventilation/perfusion matching (33). We also observed increased V_T_ in BALB/c mice compared to C57BL/6J mice, which may contribute to the higher SpO_2_ observed.

Ventilatory response data comparing 129S1 mice to C57BL/6J and CD1 mice are limited, and further research is needed to elucidate the underlying mechanism for the strain differences observed. We found apneas decreased with hypoxia exposure as we had expected, given the stimulation of the chemoreceptors by hypoxia (34). Prior studies investigating the impact of intermittent hypoxia on sleep apnea have shown that hypoxia induces respiratory plasticity and stabilizes upper airway function (35). The same mechanism may be responsible for the decrease in apneas with hypoxia exposure. However, unique to the CIH model, mice experience hypocapnia as opposed to the hypercapnia observed in humans (36, 37). This highlights the need to report changes in ventilation as V_E_/VCO_2_, as we have previously noted, particularly when making cross-strain comparisons (26).

We expected that hypoxia would induce the hypoxic ventilatory response mechanism, leading to a reduction of apneas and thereby desaturations. However, desaturations increased when mice were subjected to hypoxia. The severity of desaturations, quantified by taking the difference of average SpO_2_ during the desaturation with the average SpO_2_ of the entire F_I_O_2_ exposure period, decreased with exposure to graded hypoxia, while the duration of the events was unaffected. A possible explanation for increased desaturations during hypoxia could be motion artifact recorded by the MouseOx collar when measuring peripheral oxygen saturation, making the collar oximeter intermittently unreliable. However, mice were monitored during the experiment to ensure that measurements were only made while the mice were still, and our findings are consistent with observations made by others from our group.

Alternatively, intermittent hypoxia exposure leads to sensitization of the carotid body in mice, and this contributes to increased sympathetic tone (38). This may lead to increased peripheral vasoconstriction, causing intermittent decreases in SpO_2_. Future studies may address this by directly measuring arterial PO2 with an oxygen-sensitive probe, concurrent with the MouseOx collar oximeter measurement.

### Relationship between apneas/hypopneas and oxygen desaturation

With simultaneous plethysmography and SpO_2_ recordings, we were able to determine the relationship between desaturations and apneas. Most apneas led to desaturations, but to our surprise, desaturations were not always preceded by apneas. Only 12% of the desaturations that occurred could be attributed to apneas. To determine what may be contributing to these desaturations, we quantified hypopneas associated with desaturations. Similar to apneas, on average, only 11% of desaturations were preceded by hypopneas. While there was an increase in hypopneas leading to desaturations with decreasing F_I_O_2_, only 8% of hypopneas led to desaturations. This, along with our apnea data, suggests that mice have an alternative contributor to periodic hypoxemia. This begs the question, if not an apnea or a hypopnea, then what is the cause of periodic declines in SpO_2_? Studies on inducible intrapulmonary arteriovenous anastomoses (IPAVS), which bypass the capillary network, have demonstrated their recruitment in intact rat lungs and healthy humans during hypoxia exposure (39, 40), although it is not known whether these exist in the mouse strains studied here. Therefore, a possible explanation for these desaturations could be the result of hypoxia-induced intrapulmonary shunting. Alternatively, differences in the strength of the hypoxic vasoconstrictor response that preserves ventilation/perfusion matching may explain differences in episodic oxygen desaturations.

### Ventilatory response to graded hypoxia

Strain differences in the ventilatory response to hypoxia have been extensively studied in rats and mice (22, 23, 41, 42). The standard physiological response to hypoxia is known to be primarily an increase in breathing frequency and secondarily V_T_, or the so-called hypoxic ventilatory reflex. In our study, decreasing SpO_2_ led to an overall increase in breathing frequency (p < 0.001) but not V_T_ (p = 0.074) or V_E_ (p = 0.156). Strain was a significant predictor of V_T_, f_R_, and V_E_. For CD1, C57BL/6J, and BALB/c strain mice, the differences we observed were consistent with prior findings. For 129S1 mice, we found a novel strain difference in ventilatory responses. 129S1 mice had the greatest increase in V_T_ in response to a decrease in SpO_2_ compared to BALB/c, C57BL/6J, and CD1 mice. BALB/c and 129S1 mice had a consistent increase in f_R_ with decreasing SpO_2_. A prior study investigating the impact of hypobaric hypoxia in BALB/c and C57BL/6J mice demonstrated that BALB/c mice had a continuous increase in f_R_ while C57BL/6J mice plateaued (21). Similarly, we show that BALB/c mice had a continuous increase in f_R_ while C57BL/6J mice plateaued. BALB/c mice had the highest overall minute ventilation. 129S1 mice exhibited the greatest increase in minute ventilation in response to hypoxia, compared to other strains. This strain difference was further enhanced during the dark cycle, as indicated by a sharper increase in V_E_. Differences in the response to hypoxia may be due to adaptations to hypoxia through either reduction in metabolism or switching to ketone metabolism (23, 43). To account for changes in metabolism, we normalized minute ventilation to VCO_2_ and observed an increase in respiration in response to hypoxia (V_E_/VCO_2_). Studies investigating the impact of hypoxia on V_E_/VCO_2_ in rats have shown an increase in V_E_/VCO_2_ with respect to decreasing SpO_2_ (26). BALB/c and 129S1 mice exhibited a greater increase in V_E_/VCO_2_ in response to decreasing SpO_2_ compared to C57BL/6J and CD1 mice. Taken together, these findings underscore the importance of mouse strain in the studies of respiratory control and suggest that strain should be considered in intermittent hypoxia studies.

### Limitations

We evaluated the impact of inspired oxygen concentration on periodic desaturations and apneas in the four most commonly used strains of mice (C57BL/6J, BALB/c, 129S1, and CD1). Future studies may aim to extend these findings, particularly in the context of transgenic models. Also, the current literature lacks a consensus on a definition of what constitutes a desaturation in mice. The desaturation definition we used was based on measurements clinically relevant to humans, but the physiology of the mouse differs from that of humans, which may result in variations in the degree of SpO_2_ reduction required to classify a physiologically significant desaturation. The extent to which our findings are relevant to humans remains to be seen.

## CONCLUSION

We rigorously evaluated spontaneous apneas and desaturations in four mouse strains and found that spontaneous desaturations occur in all strains in room air. Apneas declined with reduced F_I_O_2_ but unexpectedly, desaturations increased. Surprisingly, most desaturation events were not preceded by apneas or hypopneas. Together, our findings a) provide additional evidence of strain-dependent effects on respiration; and b) suggest desaturations are frequently mediated by a mechanism outside the control of the carotid body, such as pulmonary shunting. Our data should inform investigators using mice as a model system for CIH. Future studies may elucidate the genetic and physiological basis of spontaneous desaturations in mice, and the relevance of these findings to human physiology and disease should be explored.

## DATA AVAILABILITY

Data used for this study will be made available from the corresponding author upon reasonable request.

