## Supplementary figures and images for "Spontaneous peripheral oxygen desaturation and apnea events in mice vary by strain and inspired oxygen level"

### Supplemental Figure 1

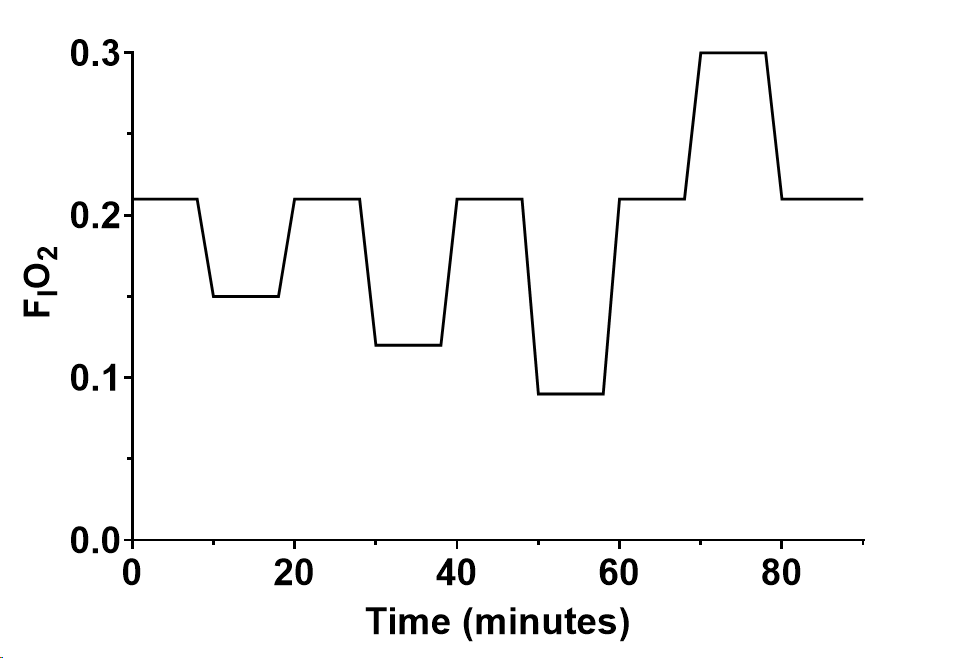
